# A yeast love triangle: multiple hybridizations shape genome evolution in the *Pichia cactophila* species complex

**DOI:** 10.1101/2023.12.27.573426

**Authors:** Verónica Mixão, Ester Saus, Ferry Hagen, Teun Boekhout, Ryan R. Stevens, Toni Gabaldón

## Abstract

Hybrids are chimeric organisms carrying genetic material from at least two divergent parental lineages. Hybridization can contribute to the emergence of novel lineages with unique phenotypic traits that may facilitate their adaptation to new environments. In recent years, genomic analyses have revealed the hybrid nature of several opportunistic human pathogenic yeasts. One of them is *Candida inconspicua,* a member of the *Pichia cactophila* species complex, for which all sequenced strains are hybrids isolated from Europe with so far unknown parentals. It has been recently proposed that *C. inconspicua* and *P. cactophila* s.s. should be ranked as the same species due to their genetic similarity. To obtain a better understanding of the evolution of this clade, we sequenced the genomes of the *P. cactophila* type strain, of its close-relative *Pichia pseudocactophila*, and of a putative *C. inconspicua* clinical isolate from Alaska and compared them with the previously sequenced genomes of *Pichia norvegensis, C. inconspicua* and the recently described *Pichia galeolata*. Our results show evidence for the existence of distinct hybrid lineages within this clade and suggest an intricate scenario of recurrent hybridizations in this species complex, some of them giving rise to lineages with the ability to infect humans. Given their different hybridization histories, we propose that *C. inconspicua, P. cactophila,* and the new clinical isolate from Alaska should represent three distinct species and suggest the name *Pichia alaskaensis* for the new taxon. Moreover, the name *C. inconspicua* is recombined in the genus *Pichia* as *P. inconspicua*. Our results clarify the evolutionary relationships within the *P. cactophila* species complex and underscore the importance of non-vertical evolution.

## Introduction

Hybridization, *i.e.* the cross of two diverged lineages, gives rise to organisms with highly heterozygous genomes, which can be a source of negative epistatic interactions, and often lead to hybrid unviability (1, 2). Different processes can shape hybrid genomes leading to loss of heterozygosity (LOH) and consequent genome stabilization (3, 4). In such cases, hybrids may survive, and their characteristic genomic plasticity may confer them some advantages, including the ability to survive in new environments (1, 5). Over the last years, a growing number of yeast hybrid lineages have been described (6), and hybridization has been hypothesized as a possible mechanism leading to the emergence of new pathogens, as it is the case of *Candida orthopsilosis, Candida metapsilosis, Candida inconspicua, Candida tropicalis* or even *Candida albicans* (1, 7–15).

*Candida inconspicua* is an emerging yeast pathogen for which all sequenced strains were isolated in Europe and correspond to hybrids with so far unknown parental lineages (7, 16). The analysis of the genomes of multiple clinical isolates revealed the existence of two distinct clades which differ in terms of genomic variability and ploidy levels. Despite these differences, it remained unclear whether the two identified clades resulted from one or two independent hybridization events (7). *C. inconspicua* belongs to the *Pichia cactophila* species complex, which also includes *P. cactophila* s.s.*, Pichia pseudocactophila Pichia norvegensis* (syn. = *Candida norvegensis*) and the recently described *Pichia galeolata*, among others (17–19) (Figure 1). While *C. inconspicua* and *P. norvegensis* are considered emerging pathogens, *P. cactophila* and *P. pseudocactophila* are more frequently found on cacti and *P. galeolata* was isolated from soil (19–24). Despite their differences in the ability to sporulate, which is almost nonexistent in the case of *C. inconspicua*, ITS-based phylogenetic analysis and identification through EF-1α sequencing and MALDI-TOF revealed a high similarity between *C. inconspicua* and *P. cactophila* (25). For this reason, the same authors suggested that these correspond to the same species (25). As a result, some studies performed since 2015 do not make any distinction between these lineages (26, 27), and propose the recombination of the name *C. inconspicua* in the genus *Pichia* as *P. cactophila* (28).

**Figure 1.**
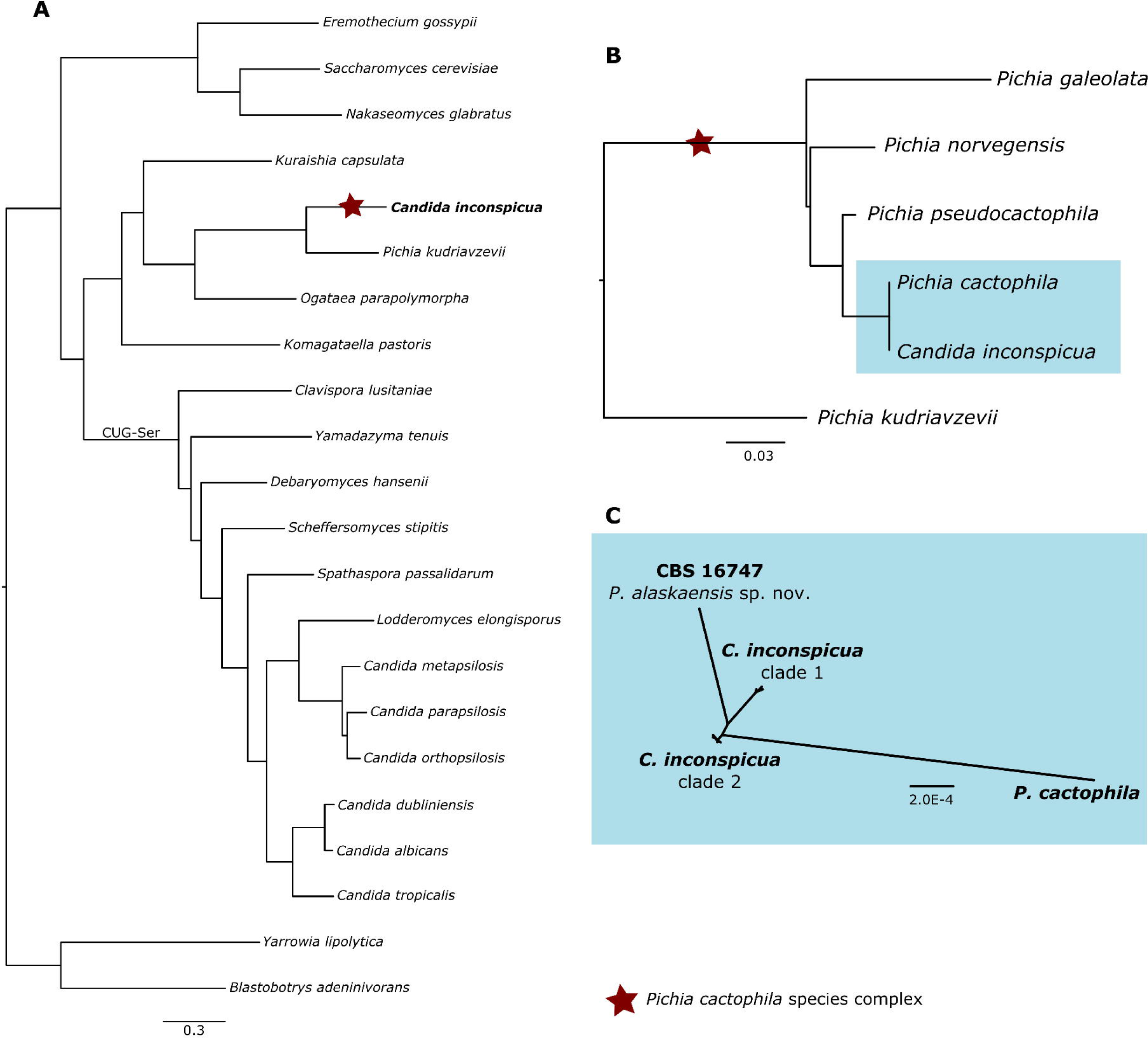
Phylogenetic analysis of the members of the *P. cactophila* species complex. **(A)** Phylogenetic tree reconstruction of the concatenated sequence alignment of the *C. inconspicua* phylome 498 (7), available at PhylomeDB (32), showing the phylogenetic placement of *P. cactophila* species complex (marked with a star) within Ascomycota. **(B)** Phylogenetic tree reconstruction of the alignment of the ITS region of the species analyzed in this study. **(C)** Phylogenetic tree reconstruction of the alignment comprising all the homozygous SNPs resultant from read mapping of *C. inconspicua* clade 1 strains, *C. inconspicua* clade 2 strains, CBS 1647 and *P. cactophila* s.s. on *C. inconspicua* reference genome (7).

Considering the hybrid nature of *C. inconspicua*, and its apparent genetic similarity to *P. cactophila*, we hypothesized that *P. cactophila* could correspond to one of its missing parental lineages. To test this, as well as to obtain further insights on the evolution of this clade, we sequenced the type strains of *P. cactophila* (CBS 6926) and *P. pseudocactophila* (CBS 6929) and performed a comparative genomics analysis with *C. inconspicua*. We have also analyzed the publicly available data of the closely related species *P. norvegensis* and *P. galeolata* (19, 29, 30), and that of a putative *C. inconspicua* clinical strain from Alaska (CBS 16747), which we propose to be ranked as a new species named *Pichia alaskaensis*.

## Materials and Methods

### Genome sequencing

Genomic DNA sequencing was performed for *P. cactophila* (CBS 6926) and *P. pseudocactophila* (CBS 6929) type-strains, and for the putative *C. inconspicua* (CBS 16747) isolated in a hospital in Alaska (USA) (31). A modified protocol from the MasterPure™ Yeast DNA Purification Kit was used to extract the DNA. In brief, samples were grown overnight in liquid YPD at 30°C. Cells were pelleted and lysed with RNAse treatment at 65°C for 15min. After 5min of cooling down on ice, samples were purified by the kit reagent by mixing, centrifugation and removal of the debris as described in the kit protocol. Further, samples were left at -20°C with absolute ethanol for at least 2h after which the DNA was precipitated for 30min at 4°C. The pellet was washed in 70% ethanol and left to dry. TE buffer was used to resuspend the DNA. Genomic DNA Clean & Concentrator kit (ZymoResearch, Irvine CA, USA) was used for the final purification.

Libraries were prepared using the NEBNext^®^ DNA Library Prep Reagent Set for Illumina^®^ kit (New England Biolabs, Ipswich, MA, USA) according to the manufacturer’s protocol. Briefly, 1 µg of gDNA was fragmented by nebulization in g-tubes (Covaris, Woburn, MA, USA) to approximately 600 bp and subjected to end repair, addition of adenine bases to the 3′-ends and ligation of Truseq adapters. All purification steps were performed using Qiagen PCR purification columns (Qiagen, Hilden, Germany). Library size selection was done with 2% low-range agarose gels. Fragments with average insert size of 700 bp were cut from the gel, and DNA was extracted using QIAquick Gel extraction kit (Qiagen) and eluted in 30 µl EB. Ten µl of adapter-ligated size-selected DNA were used for library amplification by PCR using the Truseq Illumina primers. Final libraries were analyzed using Agilent DNA 1000 chip to estimate the quantity and check size distribution and were then quantified by qPCR using the KAPA Library Quantification Kit (ref. KK4835; Kapa Biosystems, Wilmington, MA, USA) prior to amplification with Illumina’s cBot. Libraries were loaded at a concentration of 2 pM onto the flow cell and were sequenced 2 x 125bp on Illumina’s HiSeq 2500.

### Public data

For a comparative genomics analysis with *C. inconspicua*, the sequencing library of *C. inconspicua* CBS 180 (SRR8506592) and the respective genome assembly (ASM493185v1) were downloaded from NCBI database (7). Moreover, the concatenated sequence alignment of the phylome 498, which uses this strain as seed, was downloaded from PhylomeDB (32). For the analysis of *P. norvegensis* and *P. galeolata*, publicly available data was retrieved from NCBI. Specifically, we downloaded the genome assemblies under the accession numbers ASM370546v1 and ASM3055572v1, and the sequencing libraries under the sequencing runs SRR6476040 and SRR16974481, respectively (19, 29, 30).

### Read mapping and Variant calling

Next-Generation Sequencing data was inspected with FastQC v0.11.5 (http://www.bioinformatics.babraham.ac.uk/projects/fastqc/). Paired end reads were filtered for quality below 10 or size below 31 bp and for the presence of adapters with Trimmomatic v0.36 (33). Read mapping and variant calling were performed using HaploTypo pipeline v1.0.1 with default parameters (34). Briefly, BWA-MEM v0.7.15 (35) was used for read mapping on the respective genome assemblies, and GATK v4 (36) was used to call variants with the tool HaplotypeCaller followed by VariantFiltration. The read alignment was inspected with IGV v2.0.30 (37). Mapping coverage was determined with Samtools v1.9 (38). Only positions in the reference with 20 or more reads were considered for the analysis, and these were determined with bedtools genomecov v2.26.0 (39). Ploidy estimation was performed with nQuire (40).

### LOH block definition

To determine for each heterozygous strain the presence of LOH blocks, heterozygous and homozygous variants were separated. Then, the procedure applied and validated by Pryszcz and colleagues and later refined by the same group was used (8, 11). Briefly, bedtools merge v2.26.0 (39) with a distance of 100 bp was used to define heterozygous regions, and by opposite, LOH blocks would be all non-heterozygous regions in the genome (8). Moreover, the minimum LOH and heterozygous blocks sizes were established at 100 bp.

### Analysis of four universal marker genes

For the reconstruction of the phylogenetic relationships between the different hybrid lineages, the four marker genes previously proposed by Capella-Gutierrez and colleagues (41), namely, *KOG1*, *CLU1*, *VPS53* and *RFA1*, were phased in each hybrid strain using HapCUT2 (42). The different haplotypes of each gene were aligned with MAFFT v7 (43) and trimmed with trimAL v1.4.rev15 (44). RAxML v8 (45) was used to reconstruct the Maximum Likelihood phylogenetic tree of each of the multi-sequence alignments, using the GTRCAT model.

### Genome assembly

The K-mer Analysis Toolkit v2.4.1 (KAT, (46)) was used to count *k-*mer frequency and estimate the expected genome size using default parameters (*k* = 27). SOAPdenovo v2.04 (47) and SPAdes v3.9 in both SPAdes and dipSPAdes modes (48, 49) were used separately to perform the genome assembly. Afterwards, redundant contigs were removed from each assembly with Redundans v0.13c (50). The quality of the different assemblies was inspected with Quast v4.5 (51). Genome annotation was performed with Augustus v3.5 using *C. albicans* as model organism (52). The assembly completeness was estimated with KAT and BUSCO v3 using the Ascomycota database (46, 53). The best assembly for each species was chosen based on the assembly completeness, genome size, N50 and number of scaffolds.

### Phylogenetic tree reconstruction

For the phylogenetic analysis of the ITS region, the ITS sequences of *P. cactophila, P. pseudocactophila* and strain CBS 16747 were retrieved from their respective genome assemblies using the ITS sequence of *C. inconspicua* type strain (NR_111116.1) as query. Additionally, ITS sequences of *P. norvegensis, P. galeolata* and *P. kudriavzevii* were retrieved from the NCBI database (accession numbers OP800188.1, OL583853.1 and NR_131315.1, respectively). The multiple sequence alignment of these sequences was generated with MAFFT v7 (43) and trimmed with trimAL v1.4.rev15 (44).

For the whole-genome scale phylogenetic analysis of *P. cactophila* s.s., *C. inconspicua* and CBS 16747, the concatenated alignment of all the homozygous SNPs observed in at least one of these strains when mapped onto the *C. inconspicua* genome assembly (7) was used. Phylogenetic tree reconstruction was performed with RAxML v8 (45).

### Phenotypic growth test

The morphology and nutritional growth patterns of the strain of strain CBS 16747, the putative new species *P. alaskaensis*, was investigated according to standard methods used in yeast taxonomy (54). Fermentation and carbon utilization was tested in liquid media, whereas nitrogen compounds were tested on solid media using the auxanogram method. The morphology was investigated using MEA, yeast morphology agar (YMA), potato dextrose agar (PDA) and glucose yeast peptone agar (GYPA), 5% glucose in yeast nitrogen broth and Dalmau plates on yeast morphology agar. Formation of asci and ascospores was investigated for two months on the following media for which the recipes can be found in Kurtzman et al. (2011) (54): YMoA, CMA, GYPA, MEA, 1/10 YMA, Fowell acetate agar, V8-agar and McLarry agar.

### Antifungal susceptibility testing

The type-strains of *P. cactophila* (CBS 6926), *P. pseudocactophila* (CBS 6929), *C. inconspicua* (CBS 180), *P. norvegensis* (CBS 6564) and CBS 16747 (here proposed as the newly described species *Pichia alaskaensis*) were subjected to antifungal susceptibility testing according to the EUCAST broth dilution method for yeasts E.DEF 7.3.2.; http://www.eucast.org/). Antifungals tested were amphotericin B, 5-flucytosine, fluconazole, itraconazole, posaconazole, voriconazole, isavuconazole, anidulafungin and micafungin (all from Merck). The growth inhibition was checked spectrophotometrically as well as visually.

### Data availability

The clinical isolate from Alaska, which is here proposed to be ranked as a new species (*P. alaskaensis*), was deposited at the CBS yeast collection under the accession number CBS 16747. Sequencing data, and respective genome assemblies and annotation, are available at the NCBI database under the BioProject PRJNA694915.

## Results

### *P. cactophila* type strain and CBS 16747 represent two additional hybridization events in the *P. cactophila* species complex

For a better understanding of the evolution of the hybrid opportunistic pathogen *C. inconspicua*, we sequenced the type strain of *P. cactophila* (CBS 6926) and the genome of a putative *C. inconspicua* clinical isolate from Alaska (CBS 16747). Given the expected genetic proximity between these new strains and *C. inconspicua* type strain, our initial strategy was to perform a *k*-mer comparison and a read mapping approach between these new libraries and the reference genome of *C. inconspicua* (7) (details in Materials and Methods section). The *k*-mer-based analysis of CBS 16747 revealed that this strain has more than one peak of coverage, and part of the *k-*mers present in the sequencing data are absent from the *C. inconspicua* genome assembly (Supplementary Figure 1). These patterns suggest that CBS 16747 has more than one haplotype, and only one of them is represented in the assembly, suggesting that this strain may also have a hybrid origin (7). In addition, the presence of multiple peaks of coverage suggests a ploidy different from 2, and, indeed, computational estimations point that this strain has a triploid genome (nQuire histotest r^2^ = 0.98). After read-mapping and variant-calling on *C. inconspicua* genome assembly, we detected 36.39 variants/kb from which 29.04 correspond to heterozygous positions (Table 1). These levels of heterozygosity are surprisingly high for a putative *C. inconspicua* strain, when compared to previously analyzed *C. inconspicua* strains (minimum 14, and maximum 19.76 heterozygous variants/kb, (7)). Nevertheless, their distribution along the genome forming blocks of heterozygosity with similar haplotype divergence separated by blocks of LOH confirms that CBS 16747 is also a hybrid (Supplementary Figure 2). In this regard, 37.94% of this genome corresponds to LOH regions, a value consistent with a higher level of heterozygosity in this strain when compared to the previously analyzed isolates of *C. inconspicua* (minimum 51% LOH, (7)). Moreover, the current nucleotide divergence between the homeologous chromosomes of this strain is 4.46%, ∼1% higher than the previous *C. inconspicua* strains (Table 1). These differences in the levels of LOH and parental sequence divergence are unexpected for a *C. inconspicua* strain, but in accordance with the observations of the *k*-mer analysis. Altogether, these results suggest that CBS 16747 derives from a hybridization event unrelated to any of the two previously described clades of *C. inconspicua* (7).

**Table 1.**
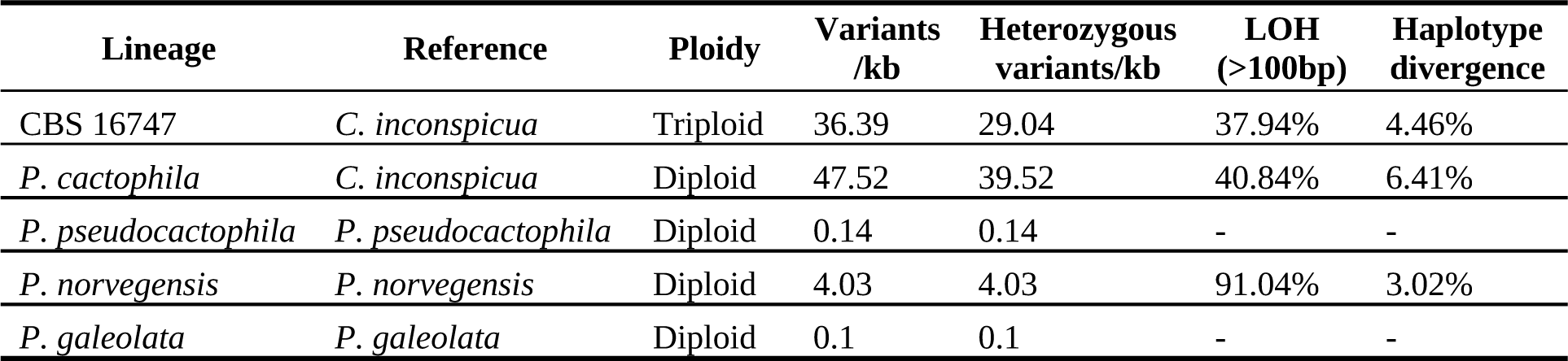
Summary of genomic variability in CBS 16747, *P. cactophila* type strain, *P. pseudocactophila* type strain, *P. norvegensis* type strain and *P. galeolata* type strain, with indication of the reference genome in which their sequencing data was aligned, ploidy, number of variants per kilo-base (kb), heterozygous variants per kb, percentage of the genome in LOH blocks, and the estimated haplotype divergence in heterozygous regions.

Similar to the analysis of CBS 16747, *P. cactophila* was analyzed based on a read mapping approach on the *C. inconspicua* reference genome. The *k*-mer comparison between these strains revealed that only part of the genome of *P. cactophila* is represented in *C. inconspicua* genome assembly (Supplementary Figure 1), and our estimations suggest that *P. cactophila* has a diploid genome (nQuire histotest r^2^ = 0.98). Indeed, this species has 47.52 variants/kb when compared to *C. inconspicua* genome assembly, from which 39.52 are heterozygous (Table 1). Interestingly, once again, the heterozygous SNPs are not homogeneously distributed along the genome. Instead, they form blocks of heterozygosity with an estimated current nucleotide divergence of 6.41% (Supplementary Figure 2), ∼3% higher than *C. inconspicua* strains (7), and ∼2% higher than CBS 16747 (Table 1), supporting a scenario of yet an additional hybridization event in the clade. Of note, *P. cactophila* has 40.84% of the genome in LOH regions. The higher sequence divergence observed in *P. cactophila* indicates that the two parental lineages of this species are not the same as the parentals of *C. inconspicua* nor CBS 16747, but the observation of shared *k*-mers between the two species suggests that they possibly share at least one of the parental lineages of their respective hybridizations. As seen in Figure 1c, the putative *C. inconspicua* strains (including CBS 16747) form different hybridization clades, and *P. cactophila* is distantly related to all of them. Importantly, these results are at odds with the previously proposed scenario that *P. cactophila* and *C. inconspicua* correspond to the same species (25).

### CBS 16747 and *C. inconspicua* share a parental lineage

To get a better understanding of the number of lineages involved in the origin of the new hybrids described here, we analyzed four previously proposed universal fungal phylogenetic marker genes (*CLU1*, *KOG1*, *RFA1*, and *VPS53*) (41) and the *MAT* locus of these strains. Briefly, we phased these genes to recover the two haplotypes in each heterozygous strain and then reconstructed their phylogenetic relationships (see Materials and Methods). Due to the absence of known parental lineages, it was not possible to concatenate haplotypes from different genes and the analysis was performed separately for each of them. The reconstruction of the different haplotypes of the four marker genes in the *P. cactophila* clade revealed at least three putative sub-clades (Supplementary Figure 3). Considering the existence of at least three different hybridization events, this suggests that some of these hybrid lineages share at least one of their parents. Indeed, for each gene and for both CBS 16747 and *P. cactophila*, one of the haplotypes is shared with *C. inconspicua* and the other one is not (Supplementary Figure 3).

In the previous study in which the genomes of multiple *C. inconspicua* strains were investigated (7), it was suggested that the polymorphisms between the two *C. inconspicua* clades observed in the *MAT* locus could be the consequence of accumulation of mutations after a shared hybridization event, and not differences between their parental lineages. These results were used to suggest that the two *C. inconspicua* clades possibly correspond to diverged lineages of the same hybridization event. However, our results show that CBS 16747 is a result of an independent hybridization event, and, even so, it shares many of the polymorphisms present in *C. inconspicua* clade 1 *MAT* **a** (Supplementary Figure 4). Thus, these polymorphisms were possibly present before both hybridization events occurred (CBS 16747 and *C. inconspicua* clade 1), and therefore *C. inconspicua* clades 1 and 2 represent two different hybridizations. Furthermore, if most of these polymorphisms are ancestral, these results suggest that the CBS 16747 and *C. inconspicua* clade 1 share the parental lineage carrying the *MAT* **a**, and differ in the donor of *MAT* **a**. This scenario is in accordance with the observation of a heterozygous ITS region in CBS 16747, with one of the haplotypes being similar to *C. inconspicua* clade 1 (Supplementary Figure 5).

### *P. cactophila* derives from a secondary hybridization involving a *C. inconspicua* hybrid and another lineage

As mentioned above, the reconstruction of the different haplotypes in the set of phylogenetic marker genes revealed that one of the haplotypes of *P. cactophila* is close to *C. inconspicua*, but the alternative one is distantly related to any of its lineages (Supplementary Figure 3). This result supports the hypothesis that *P. cactophila* and *C. inconspicua* share a parental lineage but differ in the other one. Nevertheless, further inspection of the *MAT* locus revealed a much more complex scenario. Indeed, *P. cactophila MAT* **a** seems close to that present in the *C. inconspicua* clade 1. However, the *MAT* **a** allele has a *MAT* **a1** that is different from *C. inconspicua* and CBS 16747, and a *MAT* **a2** that is heterozygous (Supplementary Figure 4). To confirm this observation, i.e., *P. cactophila* has two *MAT* **a** alleles, we assembled the genome of *P. cactophila* type strain (see Materials and Methods, and Supplementary Table 1). From this genome assembly, we recovered the full *MAT* **a** and *MAT* **a** alleles of *P. cactophila*. Our results show that one of the haplotypes of *MAT* **a2** has recombined in the *MAT* **a** and corresponds to the *MAT* **a2** of *C. inconspicua* (Figure 2a, Supplementary Figure 6). The existence of this recombination event supports a scenario in which one of the parents of *P. cactophila* was a *C. inconspicua* hybrid that underwent a recombination event in the *MAT* locus. This is in accordance with the ITS similarity observed between this species and *C. inconspicua* clade 1 (Supplementary Figure 5). It is important to note that, as mentioned before, our estimations point to a diploid state of *P. cactophila*, which is not compatible with the idea of a hybrid parent. Therefore, we suggest that, after the recombination event in the *MAT* locus, the *C. inconspicua* hybrid that gave origin to *P. cactophila* experienced a ploidy reduction before the subsequent hybridization that originated *P. cactophila* (Figure 2b). Alternatively, a *C. inconspicua* hybrid carrying the recombined *MAT* alleles crossed with the alternative parental of *P. cactophila*, and the ploidy reduction occurred afterwards.

**Figure 2.**
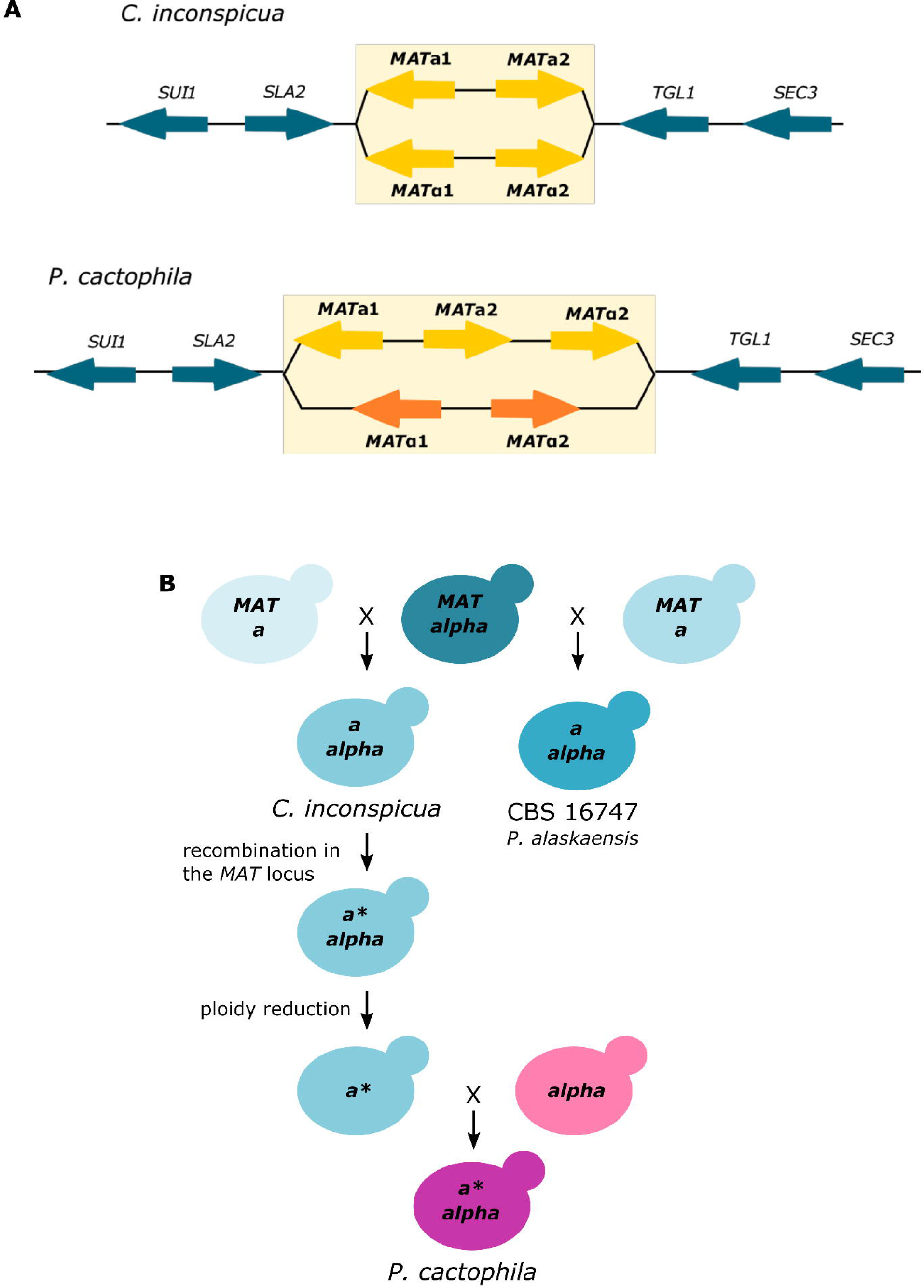
Schematic representation of *P. cactophila MAT* locus and proposed evolutionary scenario. **(A)** *MAT* locus representation of *C. inconspicua* and *P. cactophila* type strains. Yellow (*C. inconspicua*) and orange (*P. cactophila* alternative parental) arrows represent the distinct genes of the *MAT* **a** and **alpha** idiomorphs. The direction of the arrow represents the DNA strand coding the gene. *P. cactophila MAT* locus comprises two different alleles, one of them including both *MAT* **a** and a single *MAT* **alpha** gene of *C. inconspicua.* **(B)** Schematic representation of the evolutionary scenario proposed for the analyzed strains of the *P. cactophila* species complex.

### *P. norvegensis* is not a parental lineage, but rather another hybrid

With all the lineages analyzed so far in the *P. cactophila* species complex being hybrids, we decided to extend the analysis to *P. pseudocactophila, P. galeolata* and *P. norvegensis,* to assess whether one of them could be the alternative parental of *P. cactophila.* While for *P. pseudocactophila* we sequenced the type strain, for *P. galeolata* and *P. norvegensis* we retrieved the publicly available sequencing library and genome assembly from NCBI (details in the Materials and Methods section). A *k*-mer comparison showed that none of these three species is the parent of any of the hybrids identified in the clade. An inspection of *P. pseudocactophila* and *P. galeolata* revealed that both these species have a highly homozygous diploid genome (Table 1, Supplementary Figure 1, and Supplementary Table 1). In contrast, similarly to other members of the clade, *P. norvegensis* presents two peaks of *k*-mer coverage (Supplementary Figure 1), a pattern that, as mentioned before, is a good indicator of a highly heterozygous genome. *P. norvegensis* has 4.03 heterozygous variants/kb when mapped to its own reference genome (Table 1), thus presenting a lower heterozygosity than the one observed in *C. inconspicua* and *P. cactophila*. Interestingly, similarly to these other two species, the variants of *P. norvegensis* are not spread across the genome, but rather form blocks of heterozygosity, and the sequence divergence in these blocks has a single density peak (Supplementary Figure 7), indicating that the heterozygosity of *P. norvegensis* was acquired at a single time point, and that it may have a hybrid nature (8). The current sequence divergence in heterozygous blocks is 3%, and 91% of the genome is homozygous. This indicates that, despite its possible hybrid ancestry, *P. norvegensis* is much more homozygous than any other hybrid analyzed so far in the clade, pointing to an older hybridization event or to the occurrence of a massive LOH, similarly to what was previously described for the *C. albicans* clade (8, 9).

### Proposal to rank CBS 16747 as a new species: *Pichia alaskaensis*

The species concept has evolved over time and different criteria have been applied to define yeast species (55). With recent technological advances, which, for example, nowadays provide the possibility to differentiate specimens at the nucleotide level, yeast species are undergoing significant nomenclature changes aiming to correct past practices (56). In the case of the *P. cactophila* species complex, there are at least two important nomenclature uncertainties for the moment. One is the classification of *C. inconspicua* as a *Candida* species (similarly to what was in place until recently for *P. kudriavzevii*, formerly *C. krusei*, and *P. norvegensis*, formerly *Candida norvegensis*) when it is phylogenetically placed within the *Pichia* genus (Figure 1). The other one is the distinction between this same species, *C. inconspicua*, and its close relative *P. cactophila* s.s.. Indeed, although they were both initially described as different species given their morphological and phenotypic differences, the finding of their genetic similarity in the ITS, D1/D2 and large ribosomal subunit region (shown in Supplementary Figure 5) led to the ranking of both of them as the same species (25, 28). Nevertheless, the comparative genomics analysis performed in this study suggests they are different species. Although the two species are very similar in a conserved region commonly used to classify yeast species, namely the rDNA ITS region (Figure 1b, Supplementary Figure 5) (55), our results based on whole genome information clearly show that *P. cactophila* s.s. has a different genotype from *C. inconspicua* in almost 5% of its nucleotide positions. For this reason, we consider that it is important to maintain their classification as two separate species. As *C. inconspicua* does belong to the genus *Pichia,* we propose to recombine this name in the genus *Pichia* as *Pichia inconspicua* (see below).

This study uncovered another hybrid lineage within the species complex represented by strain CBS 16747. This strain was initially considered to represent *C. inconspicua* given its genetic similarity with this species in the ribosomal DNA genes, but we here showed that it does not belong to any of the previously described clades of this species (7). As shown in Supplementary Figure 5, CBS 16747 is heterozygous in the ITS region, with one ITS allele identical to that of *C. inconspicua,* likely leading to its erroneous initial classification as *C. inconspicua*, and the other ITS allele differing to the ITS of *C. inconspicua* type strain in 6 positions, suggesting it is a different species according to standard species discrimination criteria (55). The comparative genomics analysis clarifies this conundrum and shows that CBS 16747 results from the cross of one of the parents of *C. inconspicua* clade 1 and a yet unknown, more distantly related lineage (Table 1), which makes it phylogenetically distant from any of the known species of the complex (shown in Figure 1c and in Supplementary Figure 4). This genetic distance is also reflected phenotypically, as CBS 16747 presents different abilities not only regarding fermentation and metabolism of different compounds, but also regarding growth at very high temperatures and in the antifungal susceptibility profiles (Figure 3, Tables 2 and 3). Hybrid speciation is a known mechanism for the origin of novel species that is characterized by the new lineage being reproductively isolated from the two parental lineages (57). The presence of large blocks of heterozygosity across the genome indicates reproductive isolation with discrete loss of heterozygosity, as common crosses with any of the parental lineages would lead to introgression patterns characterized by the dominance of one of the haplotypes. Thus, considering genotypic, phenotypic, and phylogenetic evidence presented here we propose that CBS 16747 represents a new taxon, different from *C. inconspicua*, for which we here propose the name *Pichia alaskaensis* (see below).

**Figure 3.**
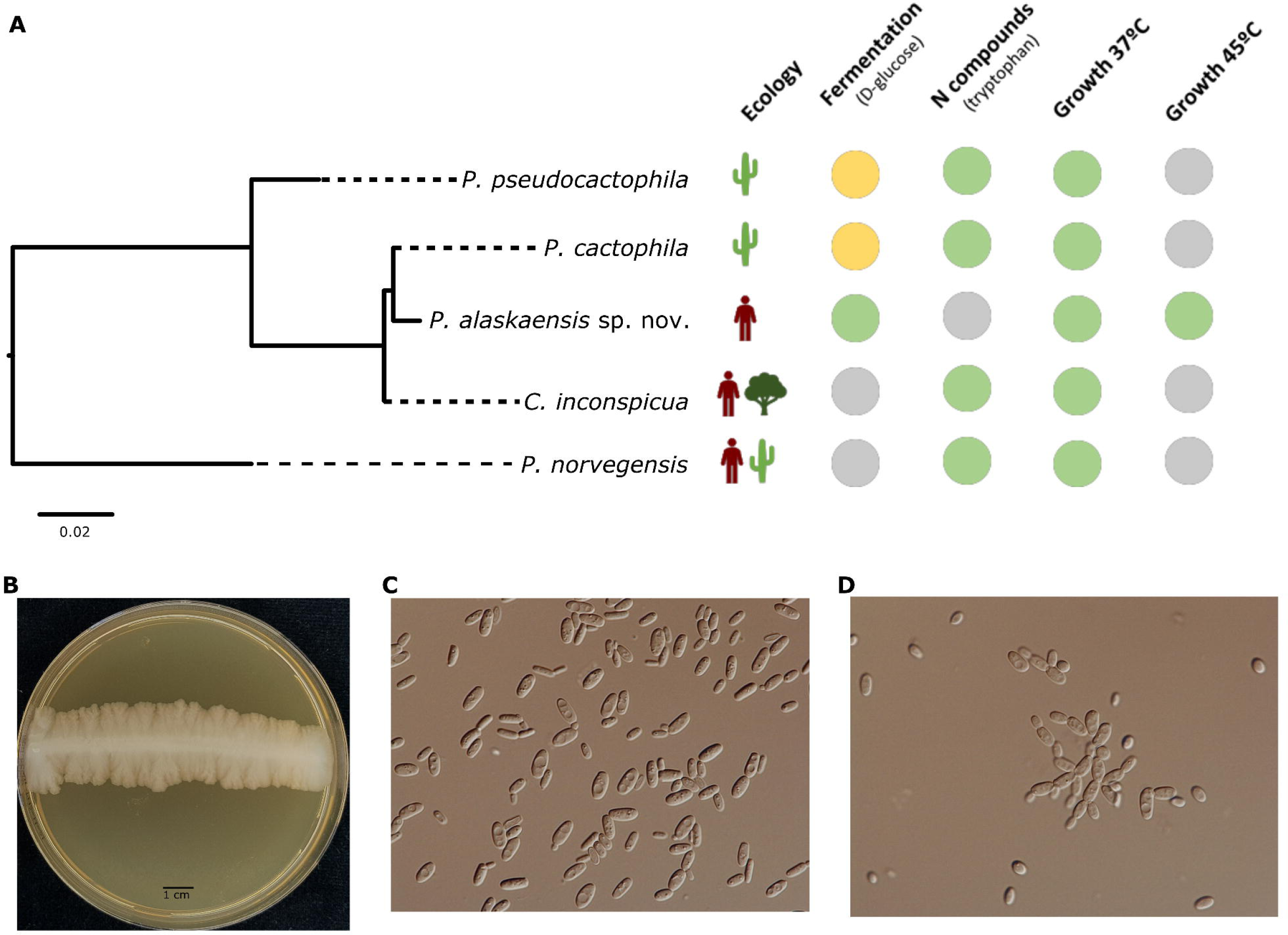
Description of *P. alaskaensis* sp. nov. (CBS 16747) genetic and phenotypic features. **(A)** Phylogenetic tree reconstruction of the nucleotide sequence alignment of the ITS region of the species of *P. cactophila* species complex analyzed in this study, with indication of their ecological niche (until the date of this study) and some phenotypic differences. Green circles indicate presence of phenotype, yellow circles indicate variable phenotype and gray circles indicate absence of phenotype. **(B)** Colonies of *P. alaskaensis* on MEA and Yeast Morphology agar (YMoA) at 25°C for 5 days. **(C)** *P. alaskaensis* cells on YYMoA at 25°C for 5 days. (D) *P. alaskaensis* cells on glucose broth at 25°C for 5 days.

**Table 2.**
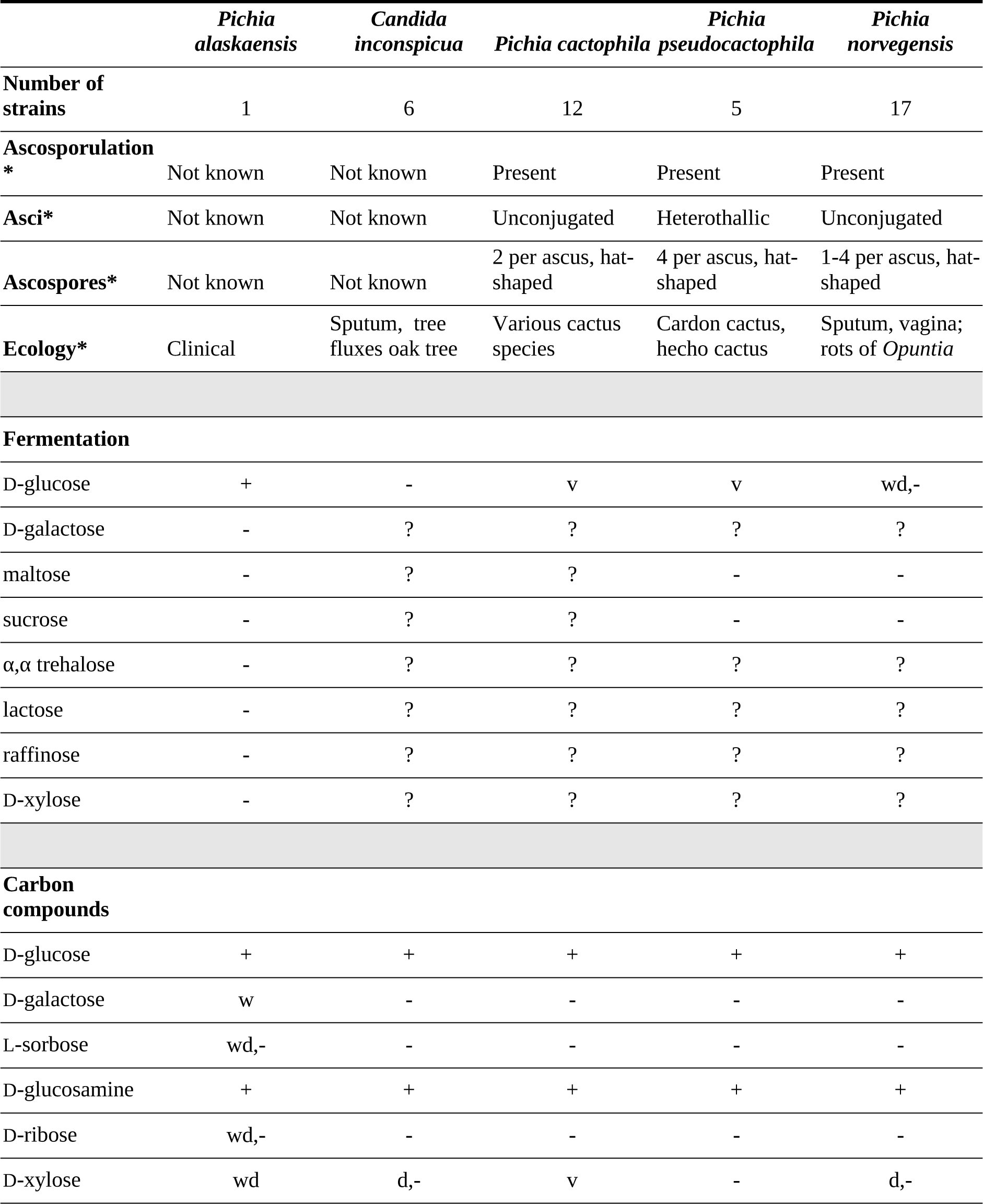
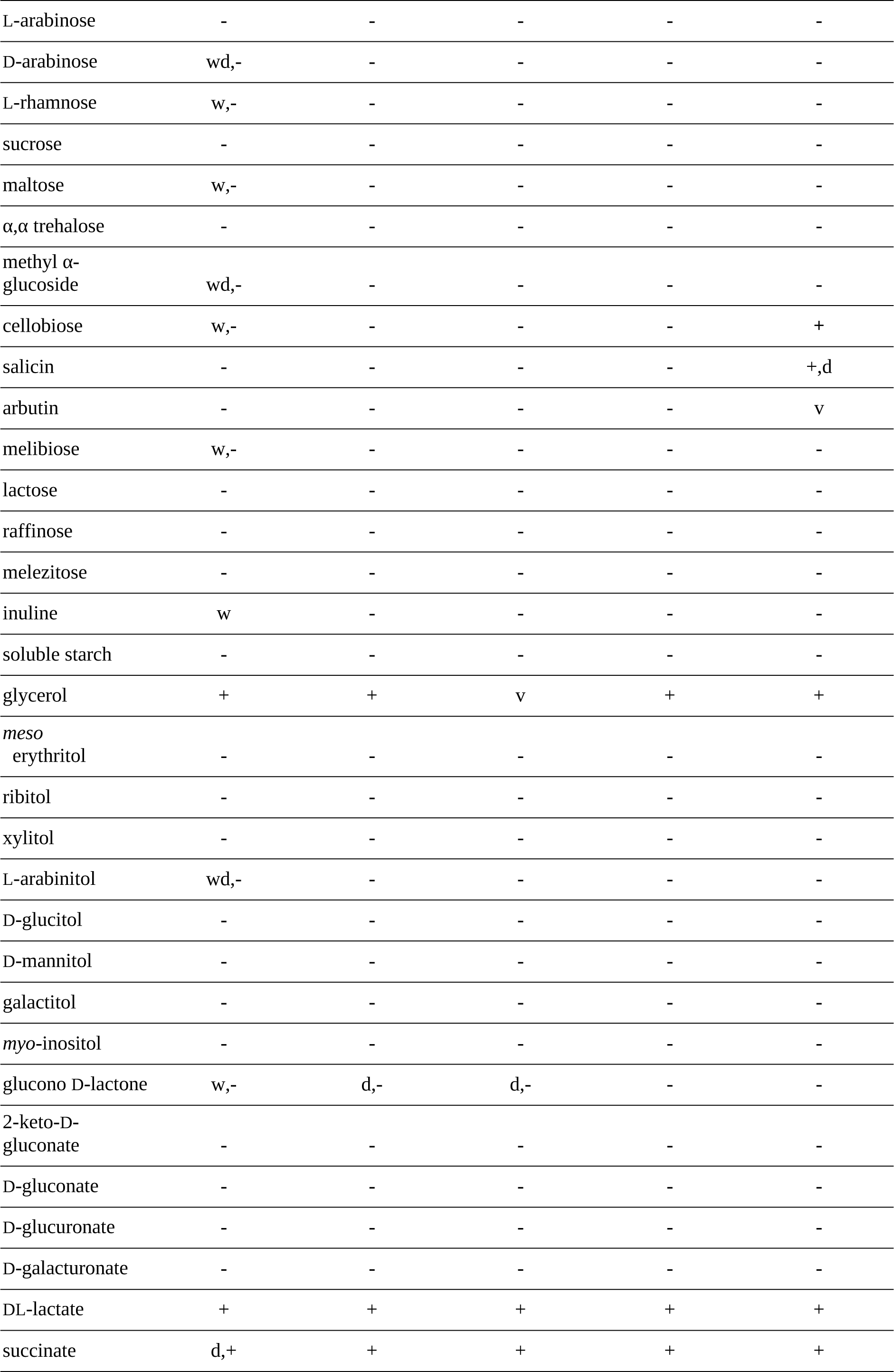
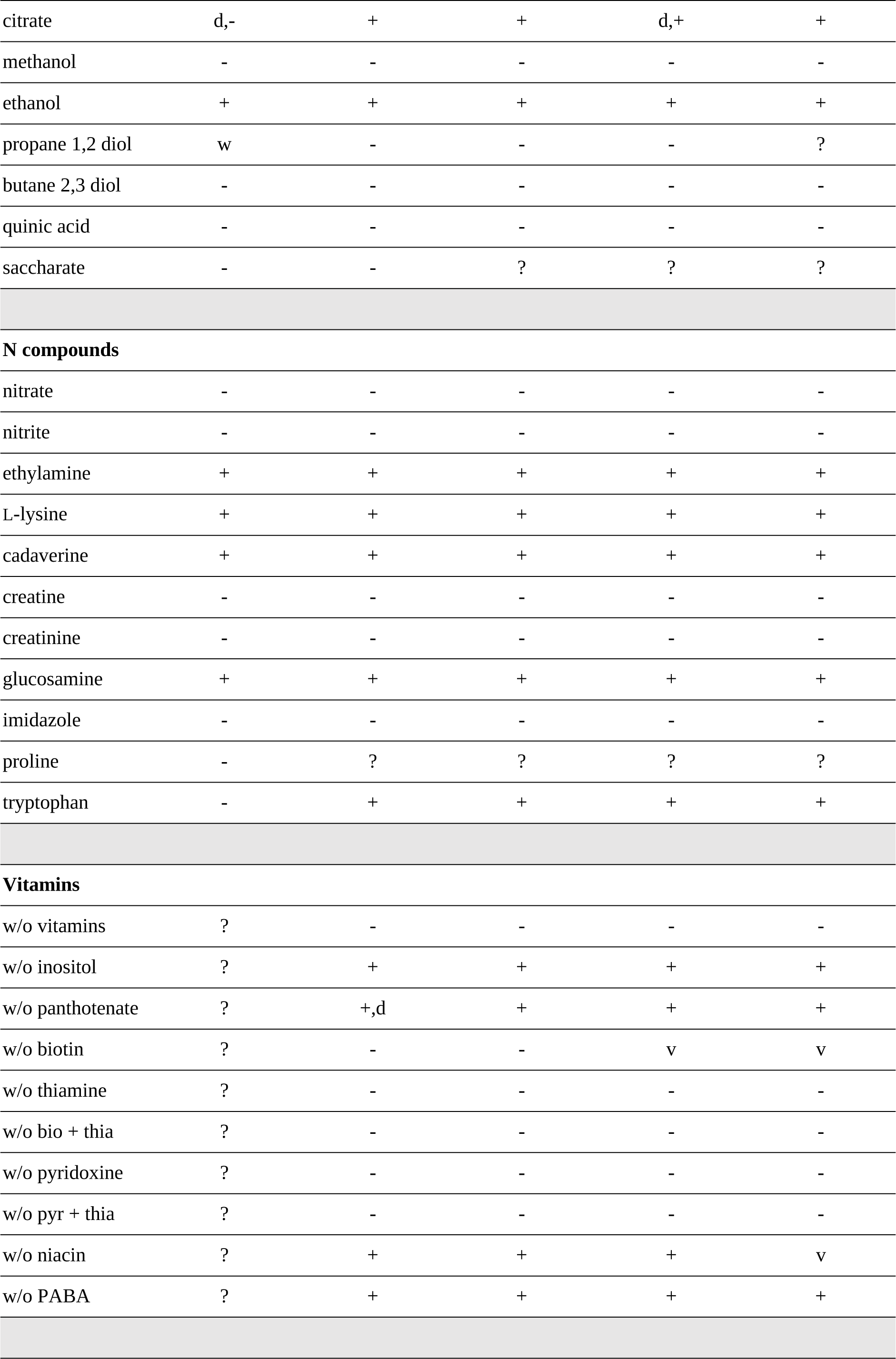
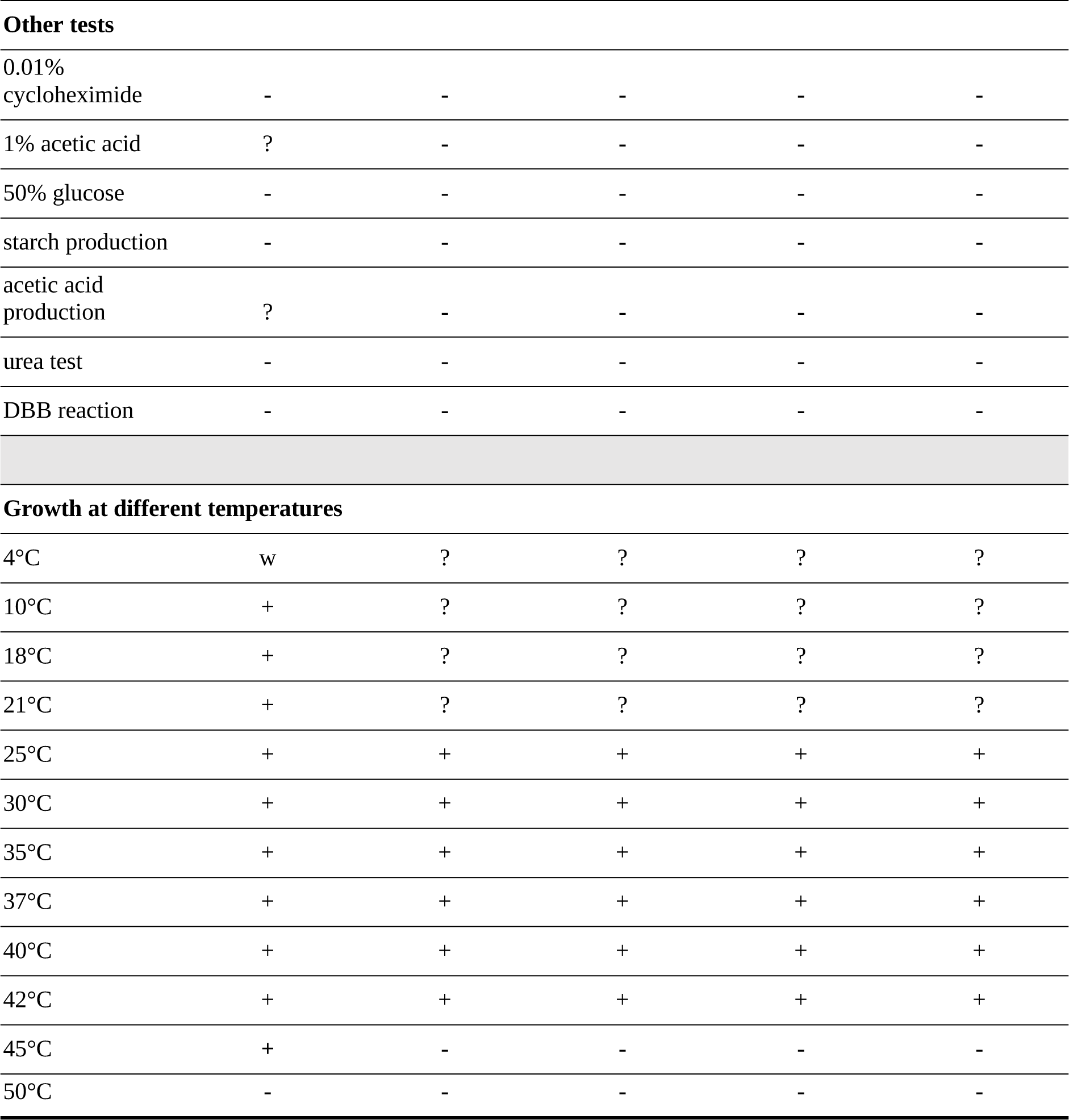
Morphological and growth features of *P. alaskaensis* and related species. Data on ecology and reproduction are taken from (64) and (65). Data on growth profiles were taken from https://wi.knaw.nl/page/Collection, except those for *P. alaskaensis* that were generated in this research, as described in the methods section. Legends: -, no growth; + growth; v, variable growth; d, delayed growth (see Kurtzman et al. 201 (64) for explanation of scores).

**Table 3.** Overview of antifungal susceptibility testing results using the EUCAST protocol.

|  | CBS 16747 | CBS 180 | CBS 6926 | CBS 6929 | CBS 6564 |
| --- | --- | --- | --- | --- | --- |
| Amphotericin B | 0.5 | 0.125 | 0.25 | 0.5 | 0.5 |
| 5-Flucytosine | 0.125 | 0.06/0.125<sup>#</sup> | 0.25 | 0.03 | 0.05 |
| Fluconazole | >64 | 64 | >64 | 32 | >64 |
| Itraconazole | 0.25 | 0.03 | 0.06/0.125<sup>#</sup> | 0.03 | 0.06 |
| Voriconazole | 0.03/0.025<sup>#</sup> | 0.03 | 0.03/0.06<sup>#</sup> | 0.03 | 0.03/0.125<sup>#</sup> |
| Posaconazole | 0.25 | 0.06 | 0.25 | 0.06 | 0.125 |
| Isavuconazole | 0.125 | 0.03 | 0.125 | 0.03 | 0.06/0.125<sup>#</sup> |
| Anidulafungin | 0.03 | 0.03 | 0.03 | 0.03 | 0.03 |
| Micafungin | 0.03 | 0.03 | 0.03 | 0.03 | 0.06/0.125<sup>#</sup> |

\* Antifungal susceptibility data is given in µg/ml; <sup>#</sup> indicates MIC-values that differ between 24h and 48h of incubation

## Discussion

In recent years, the emergence of new pathogenic lineages has shifted the epidemiology of yeast infections (16). Species that so far were not considered as medically relevant are now emerging as new opportunists (1, 16, 58). For this reason, it is important to understand their evolution. Recent studies have pointed to a role of hybridization on the emergence of new pathogens, with an increasing number of hybrid lineages being described from clinical strains (1, 6–8, 11). These comprise *C. inconspicua*, a member of the *P. cactophila* species complex, for which all clinical strains were identified as hybrids with so far unknown parental lineages (7). The absence of known parental lineages led to the hypothesis that they were possibly environmental, and hybridization possibly played a role in the emergence of this opportunistic pathogen (7). Therefore, to get a better understanding of the evolution of *C. inconspicua*, the major goal of this study was to analyze other isolates of the *P. cactophila* species complex, namely the *P. cactophila* neotype and a putative isolate of *C. inconspicua* from Alaska, USA. Our results revealed that none of these species represents the parental of the previously described hybrids, and it remains still unclear what was the role of hybridization in the emergence of *C. inconspicua* pathogenicity.

Despite our unsuccessful results on the identification of the putative parents of *C. inconspicua*, a much more complex scenario than initially thought was uncovered, involving at least four different hybrid lineages occurring in the *P. cactophila* species complex. Our results suggest that the previously described clades of *C. inconspicua* (7) correspond to two different hybridization events. This resembles the case of *Candida orthopsilosis* in which the multiple crosses of the same two lineages originated multiple pathogenic hybrids (59). More interesting, one of the parental lineages of *C. inconspicua* clades, specifically the one providing *MAT* **a**, has crossed with an alternative lineage with a higher sequence divergence than that of the alternative parent of *C. inconspicua*, giving rise to the lineage to which CBS 16747, isolated from patient from Alaska (31), belongs. Besides the possible relevance of hybridization for the ability of these lineages to cause opportunistic infections, this result raises the question of the taxonomic classification of hybrid lineages, particularly those lacking known parentals. For instance, this clinical isolate from Alaska was originally identified as *C. inconspicua* (31), but it does not result from the cross of the same two parental lineages as the *C. inconspicua* type strain. Therefore, it is uncertain whether they should have the same taxonomic classification, especially if we take into consideration their different phenotypes. This question may also be generalized to hybrids originated from a cross of the same parental lineages because the process of genome shaping may generate lineages with different genetic information, and possibly different phenotypes, despite their shared ancestry. Hybridization is a known source of genetic isolation and speciation in yeasts (6), and the case uncovered here underscores the potential complexity of resulting scenarios when the same species can originate hybrids with different partners.

The analysis of the *P. cactophila* type strain revealed that this lineage is the result of a hybridization event of a *C. inconspicua* hybrid and a yet unknown lineage. This shows that, contrary to what was previously suggested (25), *C. inconspicua* and *P. cactophila* are not the same species. The conclusion regarding the origin of *P. cactophila* was taken based on a whole-genome analysis and on a recombination event found in the *MAT* locus of this species, as it presents a recombinant *MAT* **a2** of *C. inconspicua* in the *MAT* **a** allele of the same species. Of note, such an event has previously been described in *Candida metapsilosis* hybrids (11), thus suggesting that gene conversion in the *MAT* locus may represent an important advantage for hybrid lineages. Indeed, a previous study has reported that the disruption of the *MAT* locus is important for hybrids to recover their fertility (60). Therefore, we hypothesize that this recombination event allowed a *C. inconspicua* strain to restore fertility and mate with *P. cactophila* alternative parent. Given the diploid state of *P. cactophila* we also hypothesize that after the recombination in the *MAT* locus, the *C. inconspicua* parent of *P. cactophila* restored the haploid state. Nevertheless, we cannot exclude the possibility that ploidy reduction occurred after the hybridization event giving origin to *P. cactophila.* The contribution of such a recombination in the *MAT* locus for the possible restoration of fertility and the haploid state before hybridization is still unknown. Thus, addressing the contribution of such an event for hybrids’ fertility and the emergence of new lineages is an important future step to understand their impact on genetic isolation, and, hence, speciation.

As an attempt to determine the alternative parental lineage of *P. cactophila*, we analyzed the genome of *P. norvegensis.* This analysis revealed that this species is not the parent of *P. cactophila*, and instead it represents an additional hybrid lineage. However, despite its possible hybrid nature, *P. norvegensis* has possibly undergone a massive LOH, resembling the parallel and independent massive losses of heterozygosity described for *Candida africana* and *Candida stellatoidea*, two lineages which share a hybrid ancestor with *C. albicans* (8, 9). Alternatively, if LOH is taken as a proxy for the relative age of hybrid lineages (59), *P. norvegensis* hybridization is likely much older than any of the other reported hybridization events of the *P. cactophila* species complex.

Altogether, these results show a high propensity of species of the *P. cactophila* species complex to hybridize and originate lineages with medical relevance. From the fourteen isolates of the complex which were analyzed in the past or in this study all of them are hybrids, showing the important role of hybridization on the evolution of the clade. The three new hybrid lineages here described are part of a growing number of yeast hybrids that are described using next-generation sequencing data, even in the absence of known parental lineages (1, 6–9, 11, 12, 61). This is only possible due to the characteristic genomic patterns of hybrid genomes, which are only identified when looking at the whole genome level, and not at a single gene. For instance, the analysis of ITS showed a high genetic similarity between *C. inconspicua* and *P. cactophila*, leading to the proposal that they should be considered as the same species (25, 28). Although the species concept is difficult to be applied in a context of non-vertical evolution, we consider of extreme relevance the distinction between these lineages, as they present different phenotypic traits (Figure 3 and Table 2). Indeed, both lineages were thought to be associated with different environments (20–24), and perhaps this is the reality. However, if a similar species name is used, we will lose the power to distinguish their geographical and host distribution. A similar scenario is found for the CBS 16747 strain, which due to its high similarity to *C. inconspicua* in highly conserved genetic regions, such as ITS, was classified as a member of this species. Nevertheless, both the genomic analysis and the phenotypic assays performed in this study showed that this isolate is significantly different from *C. inconspicua,* which led us to propose its ranking as a new species named *Pichia alaskaensis*. The distinction between the species of this complex is particularly relevant in the clinical setting, as for instance it would be relevant to distinguish opportunistic infections caused by *C. inconspicua* and *P. cactophila* for the assessment of their potential as emerging pathogens. Therefore, there is an urgent need for proper guidelines for lineage classification in a context of hybridization.

## Description of *Pichia alaskaensis* sp. nov

### *Pichia alaskaensis* Gabaldón, Mixão, Hagen, Boekhout, spec. nov

Mycobank number: MB 851200

Etymology: alaskaensis, from Alaska, the state, USA, where the species was found

## Morphology

Growth in glucose fermentation broth with a thin whitish film, somewhat adhering to the glass of the test tube, and with white sediment.

Colony on yeast morphology agar after 7 days at 25°C is 6-10 mm width, flat, smooth, shiny, off-white, butyrous, reverse whitish, with straight and entire margin (Figure 3). On malt extract agar, it is 17 mm width, somewhat striate near margin; Dalmau culture on malt extract agar and potato dextrose agar showed patches of pseudohyphae.

Yeast cells in glucose fermentation broth are ellipsoidal to cylindrical, 6.0-8.5 × 3.0-5.0 μm, with polar, sympodial to multilateral budding with the buds sessile on a rather broad base, single, in pairs or in short, branched chains that from small clusters of cells. Yeast cells on yeast morphology agar and malt extract agar similar, 6.0-10.5 × 2.8-5.5 μm, with cylindrical cells forming pseudohyphae that form laterally blastoconidia. After two months at YMoA larger cells occur, 8.0-11.0 × 2.0-5.0 μm, with polar and sympodial budding and that adhere into short pseudohyphae. No sexual state has been observed on YMoA, CMA, GYPA, MEA, 1/10 YMA, Fowell acetate agar, V8-agar and McLarry agar after up to two months of observations.

For the growth profile of *P. alaskaensis*, see Table 2. Glucose is fermented, carbon utilization is similar to that of the related species and thus does not distinguish the species. The same holds for the utilization of nitrogen compounds. Contrary to the related species, *P. alaskaensis* can grow at 45°C.

The MIC values of *P. alaskaensis* for the antifungal drugs tested are presented in Table 3. For all antifungal drugs tested, the strain yielded low MIC values, except for fluconazole to which *Pichia* species are intrinsically resistant (62). The MIC-values for amphotericin B, itraconazole, voriconazole, posaconazole, anidulafungin and micafungin for CBS 16747 were below that of the epidemiological cut-off values (ECOFFs) for the clinically relevant species *P. kudriavzevii* (=*Candida krusei*) as outlined by Astvad et al (2022) (63). For 5-flucytosine and isavuconazole there are no ECOFFs defined.

### Origin of strain(s)

#### Holotype

CBS 16747, preserved metabolically inactive in the Westerdijk Fungal Biodiversity Institute, Utrecht, the Netherlands. CBS 16747 was, in 2018, isolated from the blood of a patient in Anchorage, Alaska, USA

Ex-type isolates: TG20210326 (collection Toni Gabaldón) = 2MG-A1202-16 (working collection Medical Mycology research group at WI).

Note: *Pichia alaskaensis* can be phenotypically distinguished from its close relatives by its ability to grow at 45°C, to grow with D-galactose and inuline as sole carbon sources, lack of growth with D-tryptophane as a nitrogen source, and higher MIC values for itraconazole (Tables 2 and 3).

#### New combination

*Pichia inconspicua* (Lodder & Kreger-van Rij) Gabaldón, Mixão, Hagen, Boekhout, comb. nov. Mycobank number: MB 851201

Basionym: *Torulopsis inconspicua* Lodder & Kreger-van Rij, 1952. The Yeasts: a Taxonomic Study: 671 (1952) [MB#306936]

Obligate synonym: *Candida inconspicua* (Lodder & Kreger-van Rij) S.A. Meyer & Yarrow (1978) [MB#310280]

Facultative synonym: *Torulopsis inconspicua* Lodder & Kreger-van Rij variety *filiforme* Dietrichson (1954). [MB#347647]

Note: In Mycobank *Candida inconspicua* variety *thermotolerans* O.G. Lima, M.H. Maia & F.C. Albuq. is listed under Mycobank # 350416, but according to this website this variety was not validly described as no Latin diagnosis was provided. Also, no strain is known to be preserved, thus its identity remains not clear.

## Declarations

### Availability of data and material

Data generated during this work, as whole-genome sequencing data and *P. cactophila* genome assembly can be found in the NCBI database under the BioProject PRJNA694915.

## Acknowledgments

We would like to thank Elaine Francisco for generating the antifungal susceptibility data, and Dr. Benjamin Westley for his help in coordinating shipment of the clinical strain.

## Supplementary material

**Supplementary Table 1.** Summary statistics of the genome assemblies of *P. cactophila* type strain, *P. pseudocactophila* type strain and *P. alaskaensis* sp. nov. (CBS 16747).

**Supplementary Figure 1.** *K*-mer plots of the newly sequenced isolates, indicating the number of different *k*-mers across the genome coverage. Red represents the *k*-mers also observed in *C. inconspicua* reference genome and black represents the absent ones. **A)** CBS 16747 (*P. alaskaensis* sp. nov); **B)** *P. cactophila*; **C)** *P. pseudocactophila*; **D)** *P. geolata*; **E)** *P. norvegensis*.

**Supplementary Figure 2.** Density of the number of heterozygous SNPs per 100 bp in the heterozygous blocks determined for CBS 16747 (*P. alaskaensis* sp. nov) and *P. cactophila* type strain when aligned to *C. inconspicua* genome assembly.

**Supplementary Figure 3.** Phylogenetic tree reconstruction of the alignment of the KOG1 **(A)**, CLU1 **(B)**, RFA1 **(C)** and *VPS53* **(D)** phased haplotypes of *C. inconspicua* strains (7), CBS 16747 (*P. alaskaensis* sp. nov.) and *P. cactophila* type strain. For clearness reasons, CBS 180 is the only *C. inconspicua* strain indicated.

**Supplementary Figure 4.** IGV screenshot of the alignment of the library of *C. inconspicua* clade 2 (represented by the type strain CBS 180), *C. inconspicua* clade 1 (represented by 14ANR23920), CBS 16747 (*P. alaskaensis* sp. nov.) and *P. cactophila* on the *MAT* **a (A)** and *MAT* **a (B)** regions of *C. inconspicua* reference genome. In this figure, it can be seen the similarity of CBS 16747 *MAT* **a** to that of *C. inconspicua* clade 1 and their differences in *MAT* **a**. This figure also shows the differences between *P. cactophila* and *C. inconspicua* in the *MAT* **a** and their similarities in *MAT* **a**. The recombination event in the *MAT* **a** of *P. cactophila* is highlighted in red.

**Supplementary Figure 5.** IGV screenshot of the alignment of the library of *C. inconspicua* clade 2 (represented by the type strain CBS 180), *C. inconspicua* clade 1 (represented by 14ANR23920), CBS 16747 (*P. alaskaensis* sp. nov.), *P. cactophila* and *P. pseudocactophila* on the region that goes from the 18S small subunit ribosomal RNA gene (partial sequence) until the 26S large subunit ribosomal RNA gene, partial sequence.

**Supplementary Figure 6.** Screenshot of a portion of the sequence alignment of *P. cactophila MAT* a that comprises the recombinant site between *MAT* **a2** and *MAT* **a2** (PICCA_alpha2_a2) and the *MAT* **a2** (CANINC_a2) and **a2** (CANINC_alpha2) sequences of *C. inconspicua.* The recombinant site is highlighted in pink.

**Supplementary Figure 7.** Genomic patterns of *P. norvegensis* and *P. pseudocactophila*. A) IGV screenshot of *P. norvegensis* genome showing the occurrence of heterozygous polymorphisms separated by what possibly are blocks of LOH. **B)** Density of the number of heterozygous SNPs per 100 bp in the heterozygous blocks determined for *P. norvegensis* when aligned to its own genome assembly.

