## Supplementary figures and images for "A yeast love triangle: multiple hybridizations shape genome evolution in the *Pichia cactophila* species complex"

### Supplementary_Figure_1.tif

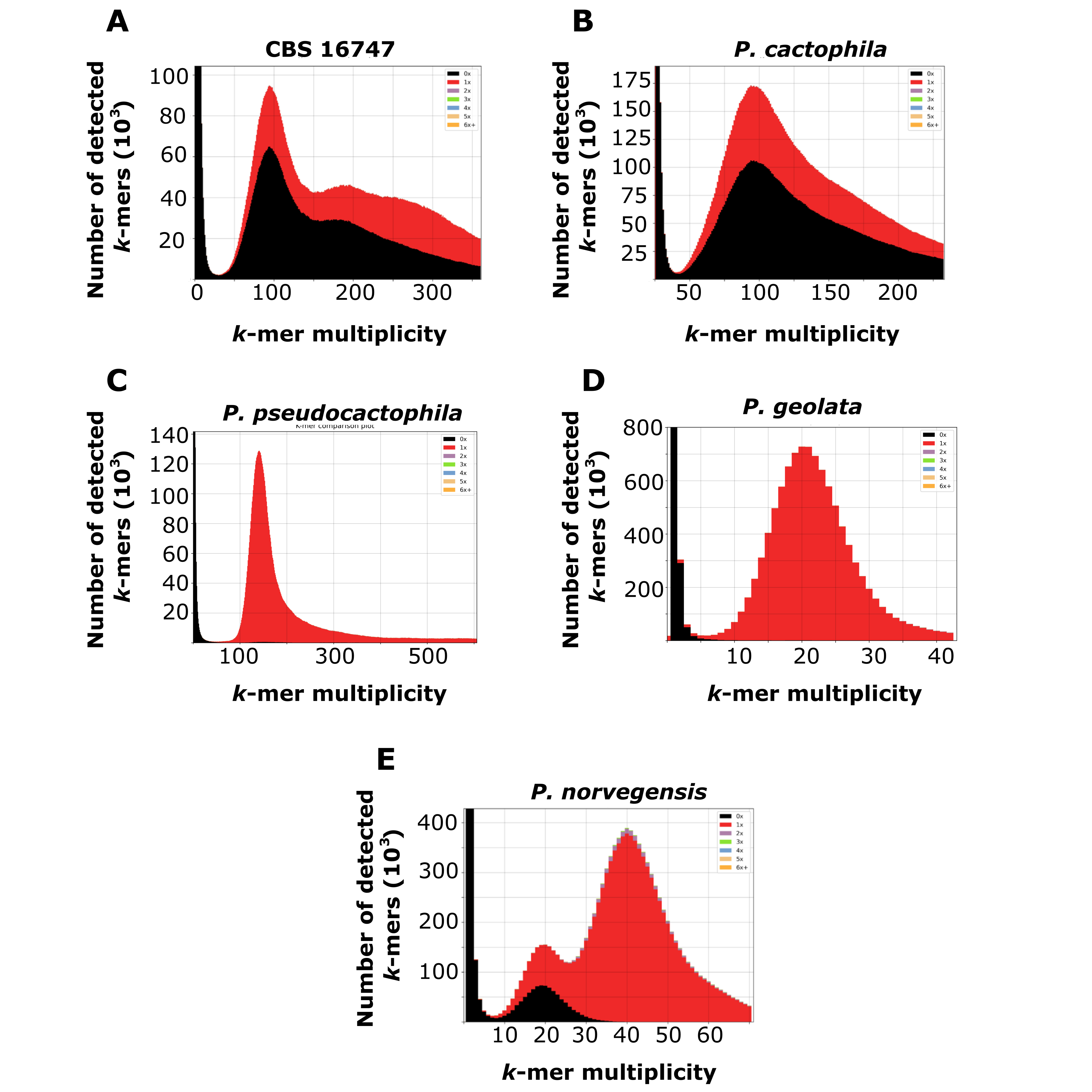

### Supplementary_Figure_2.tif

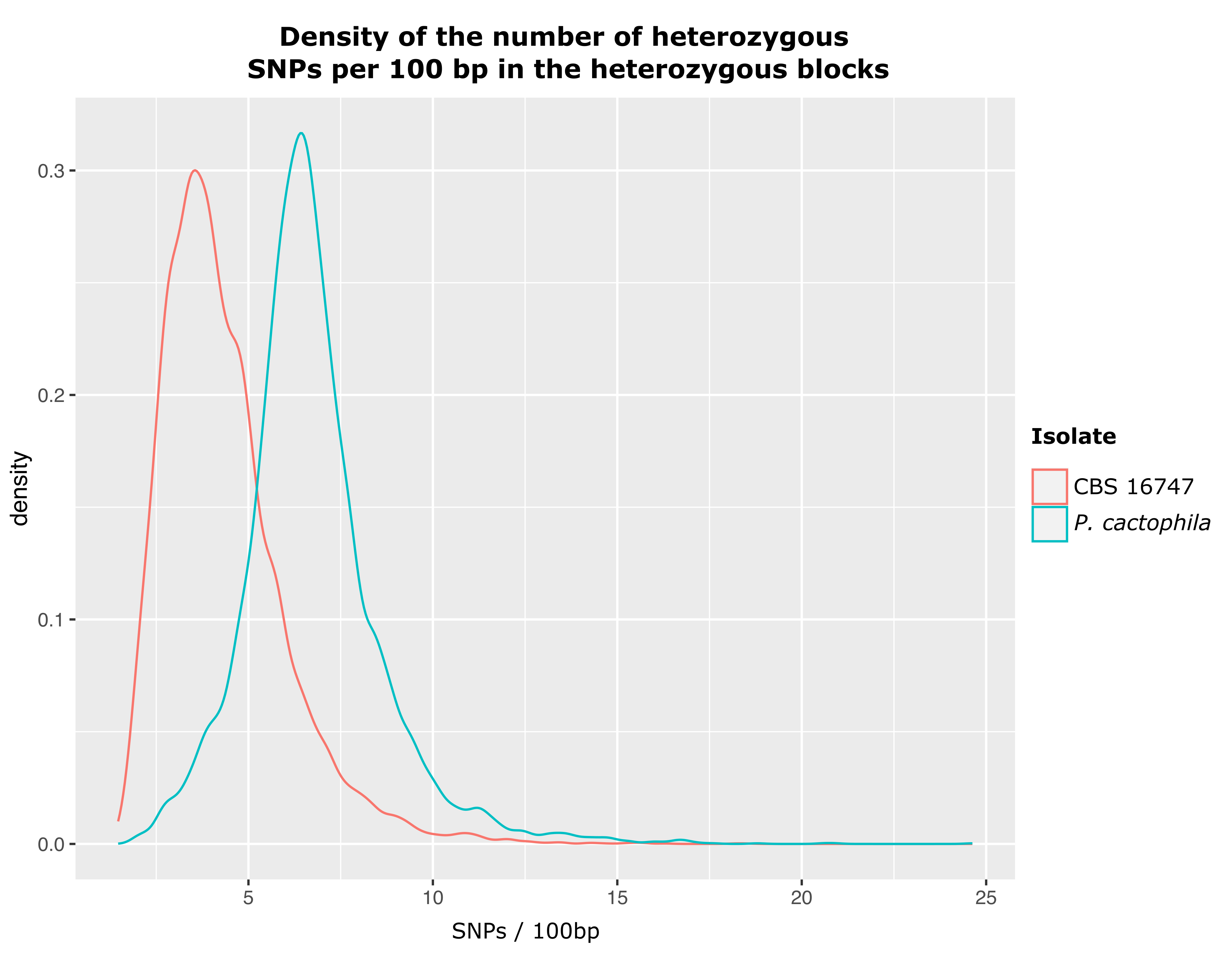

### Supplementary_Figure_3.tif

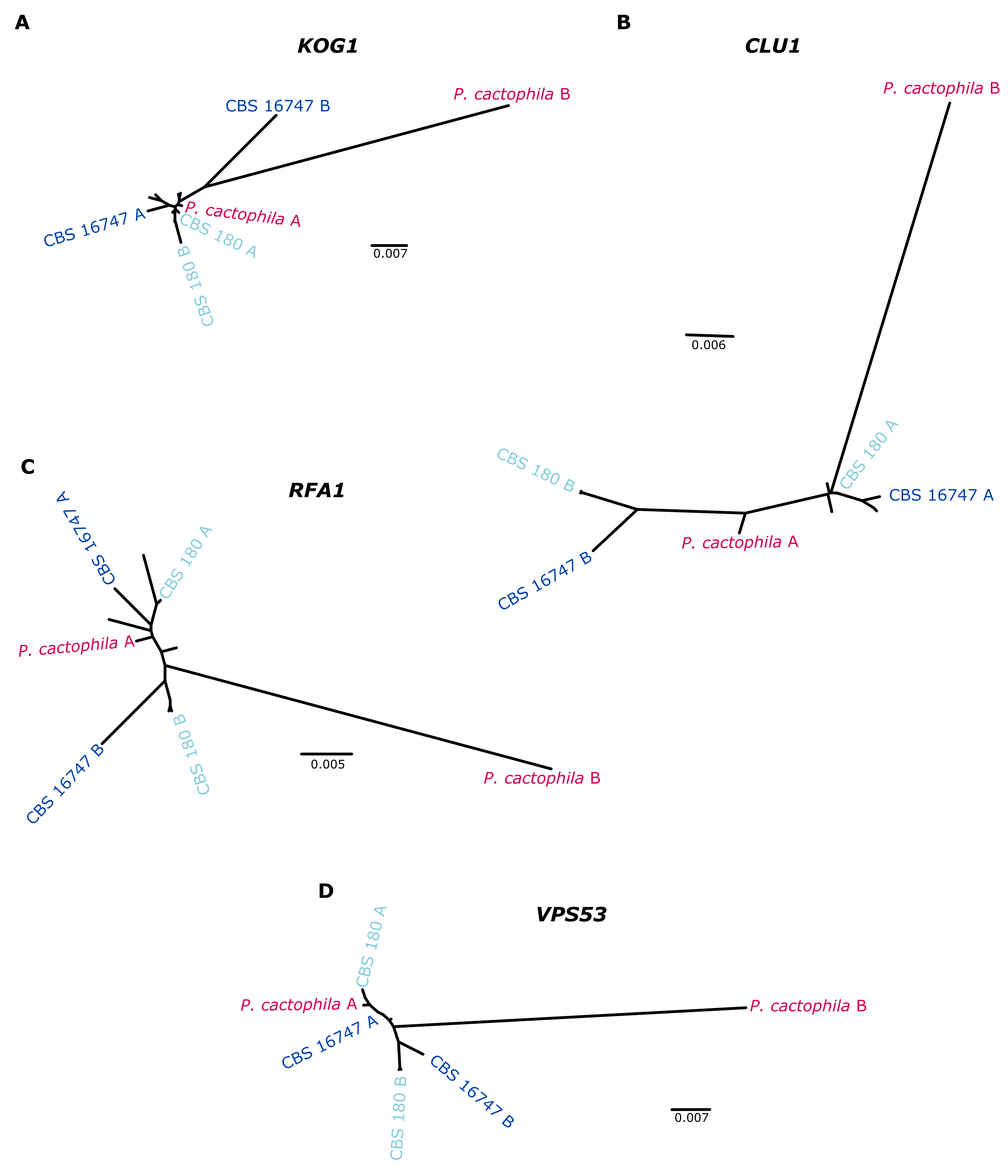

### Supplementary_Figure_4.tif

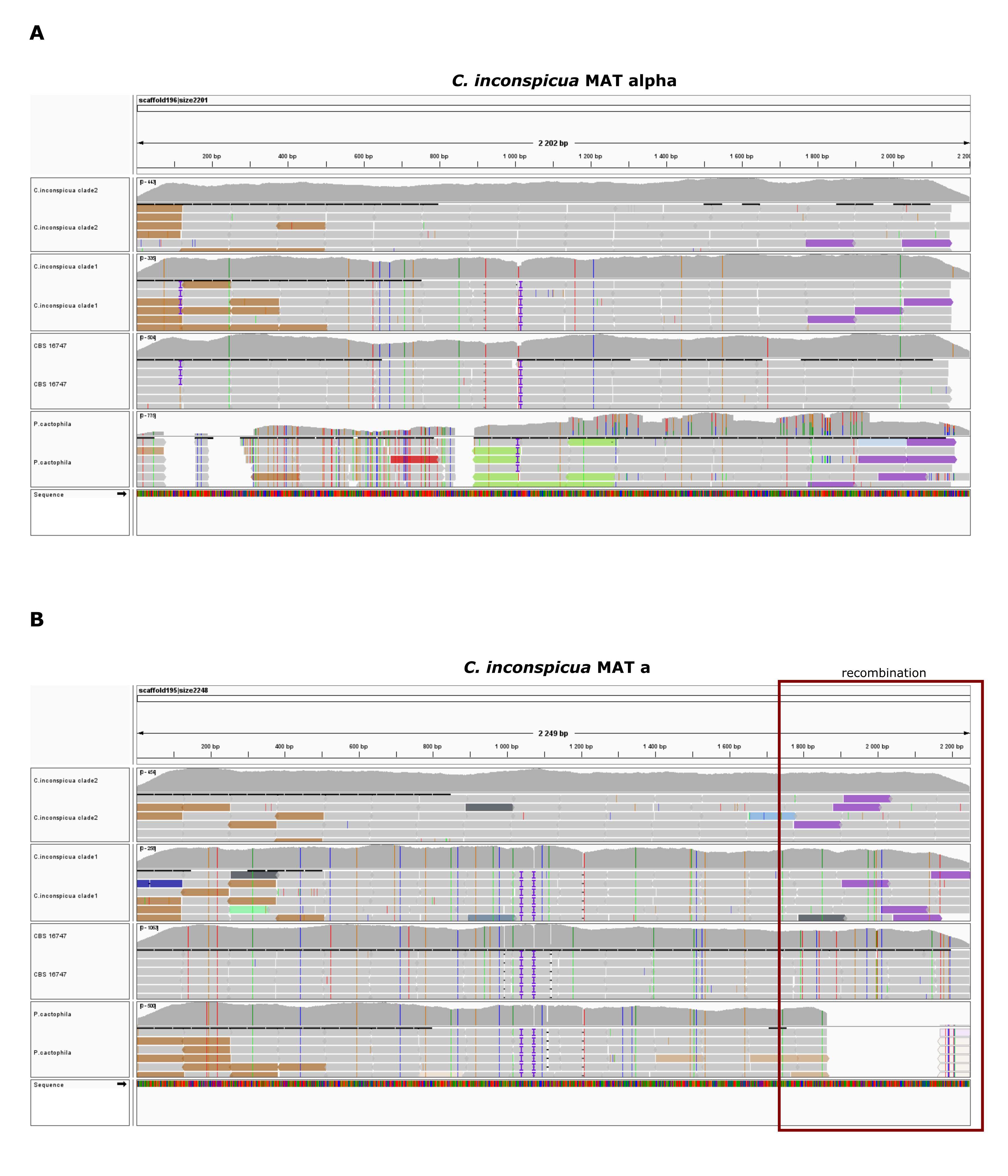

### Supplementary_Figure_5.tif

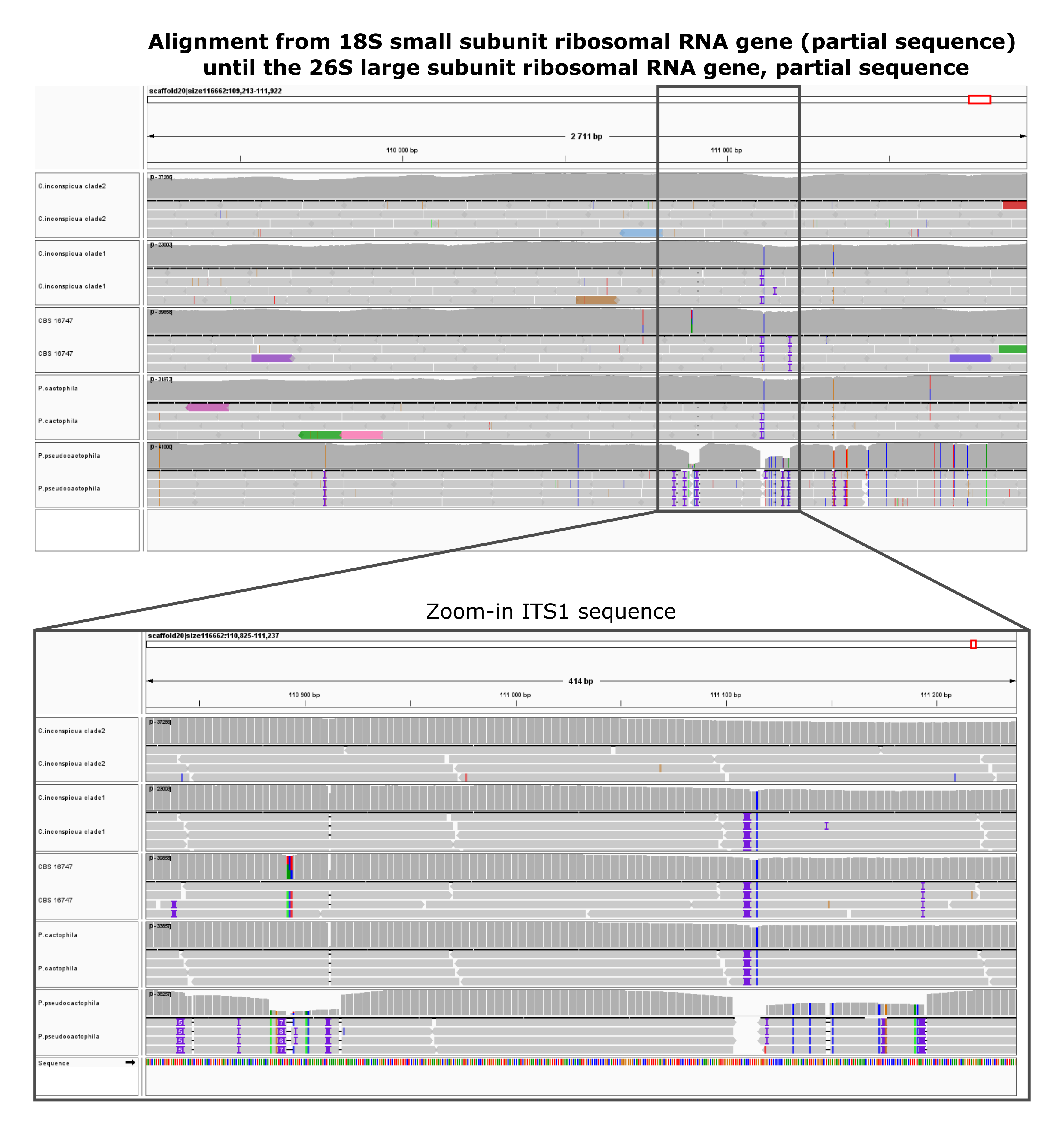

### Supplementary_Figure_6.tif

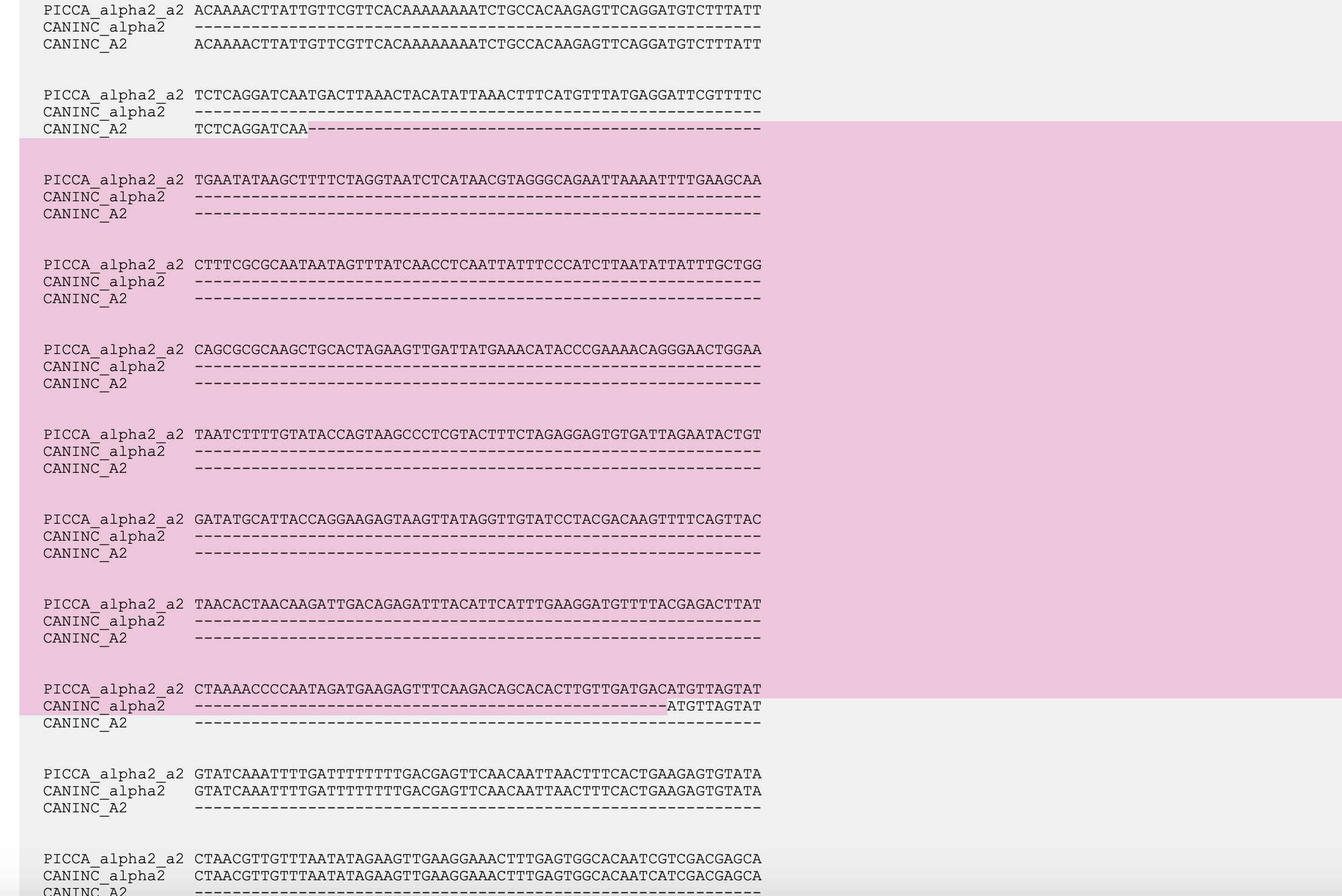

### Supplementary_Figure_7.tif

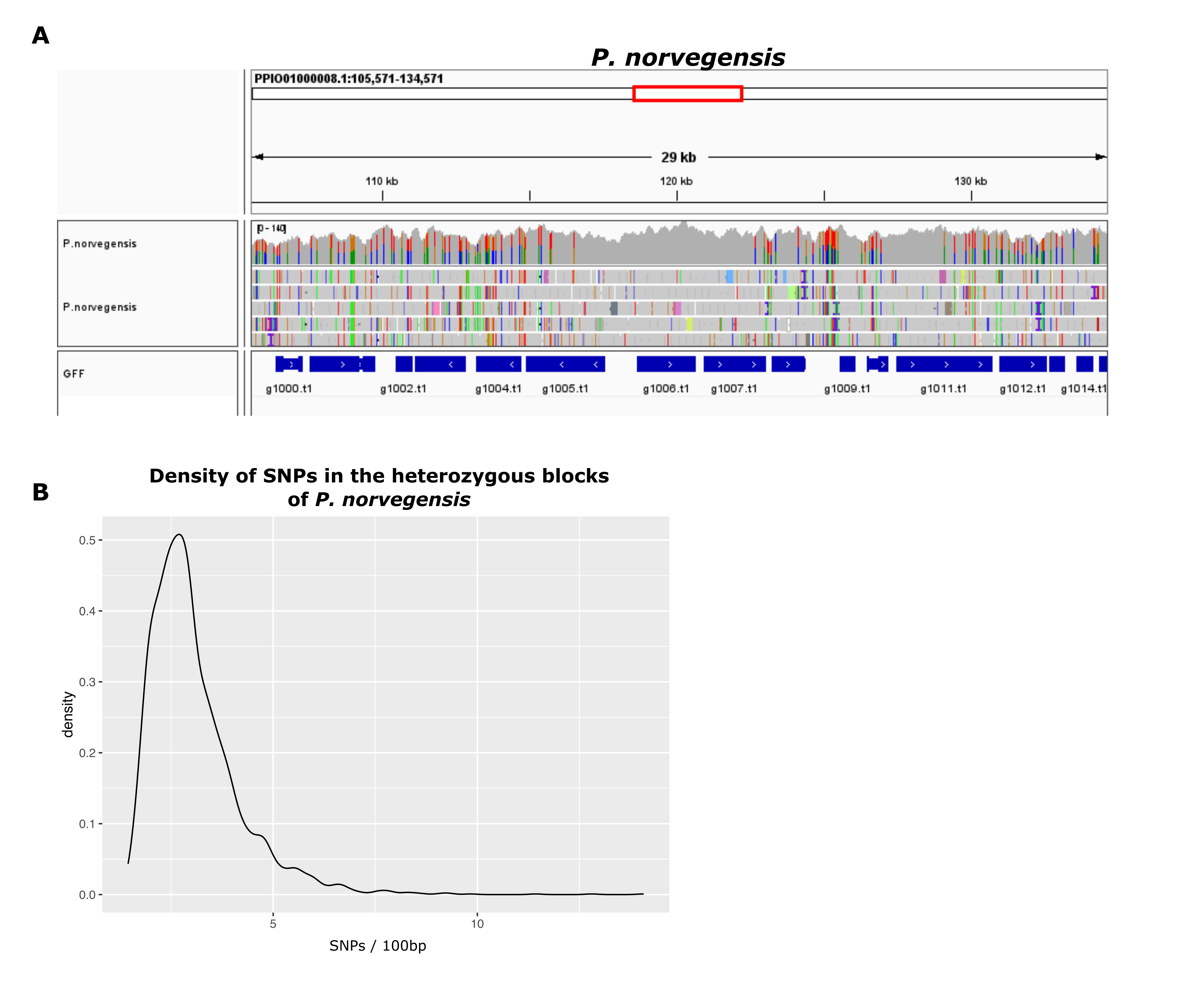
